# Scope+: An open source generalizable architecture for single-cell atlases at sample and cell levels

**DOI:** 10.1101/2022.12.03.518997

**Authors:** Danqing Yin, Yue Cao, Junyi Chen, Candice L.Y. Mak, Ken H.O. Yu, Yingxin Lin, Jiaxuan Zhang, Jia Li, Joshua W. K. Ho, Jean Y.H. Yang

**Author notes:** equal contribution (Danqing Yin, Yue Cao, Junyi Chen). equal contribution (Yingxin Lin, Joshua W. K. Ho, Jean Y.H. Yang). Correspondence: Prof Jean Yang.

## Abstract

With the recent advancement in single-cell technologies and the increased availability of integrative tools, challenges arise in easy and fast access to large collections of cell atlas. Existing cell atlas portals rarely are open sourced and adaptable, and do not support meta-analysis at cell level. Here, we present an open source, highly optimised and scalable architecture, named Scope+, to allow quick access, meta-analysis and cell-level selection of the atlas data. We applied this architecture to our well-curated 5 million Covid-19 blood and immune cells, as a portal, Covidscope (https://covidsc.d24h.hk/). We achieved efficient access to atlas-scale data via three strategies, such as server-side rendering, novel database optimization strategies and an innovative architectural design. Scope+ serves as an open source architecture for researchers to build on with their own atlas, and demonstrated its capability in the Covidscope portal for an effective meta-analysis to atlas data at cellular resolution for reproducible research.

## Introduction

Single-cell technologies have become one of the most powerful and popular techniques for understanding cell identity and function at the individual cell level. With the maturity and accessibility of the technologies, there has been a surge of interest in utilising single-cell technologies on profiling cells from large human cohort ^1,2^ to investigate the cellular response and mechanisms at population level, which is essential to discover how cellular heterogeneity contributes to an individual’s disease outcomes. This includes the study of the COVID-19 pandemic, which has had a profound influence on human life worldwide. Since the start of the outbreak, researchers from around the world have been collecting single-cell data from COVID-19 patients with diverse characteristics, degrees of severity, and disease outcomes to understand the cellular response of the disease. This includes the study of peripheral blood immune responses and mechanisms in COVID-19 patients at an atlas-scale ^3,4^, the study of epithelial-immune cell interaction and antiviral innate immunity in the upper airways ^5–7^, and the investigation of immune response variation across patients with diverse severity and outcomes ^8,9^.

Despite having a myriad of single-cell data sets, it remains a challenge to effectively access, organise, query, share and analyse multiple data sets jointly. Recently, some of these efforts resulted in the creation of a number of atlases - collating single-cell omics data from multiple studies. These include large scale atlases for healthy and diseased organs in the Human Cell Atlas ^10–13^, human cells ^14,15^; cancer tumour cells ^16–18^, see Supplementary Table 1 for more details. In the context of COVID-19, there is also an increasing number of curated data sets with varied degrees of curation. Sungnak and colleagues ^19^ provide an initial repository of study-specific access to various COVID-19-related studies in the Human Cell Atlas; and to various dashboards providing integrated or merged data sets of varying sizes: 0.48 million cells ^1^, and 1 million cells ^20^. One of the most current compilations of COVID-19 single-cell data set consists of 2.5 million peripheral blood mononuclear cells (PBMC), with an associated portal that offers exploratory visualisation and HDF5 file download of the combined data set ^21^. A similar size COVID-19 collection is found in DISCO, with 2.6 million COVID-19 cells.

While these atlases contain much information, each differs in its ways of storing and users’ accessing data. This access ranges from the degree of raw data access to the level of processing. This includes simple browsing of sample metadata with filters ^10,22^ or repositories for simple visualisations and access to analytical results ^1,14,19,21^; as well as combinations of all these ^13,15–18,20^. More recently, more sophisticated access has been made available, allowing subsetting of the single-cell gene expression data based on cell barcode or sample standardised metadata ^15^ and downloading for further analysis. However, a survey on existing single-cell atlas portals shows that they are not open sourced, and do not release as a generalizable architecture for adaptation and they are unable to subset the data at cell-level for selective downstream analysis, as compared to our Scope+ in the table (Supplementary Table 1).

The large-scale single-cell RNA-sequencing (scRNA-seq) data brought by the HCA and other initiatives poses numerous challenges in assembling data from different sources into a comprehensive atlas ^15^. A fundamental challenge is due to the natural characteristics of single-cell data, where the number of genetic features (i.e., rows) is typically over 20,000 and the number of single cells in a single study (i.e., columns or records) is generally in the range of hundreds of thousands. Traditional relational databases, which usually have limitations on the number of dimensions, are incapable of storing data at the atlas level of magnitude. Additionally, dedicated infrastructure is required to provide ensemble storage and fast querying of massive single-cell multi-omics data. Importantly, while conventional atlases support upload of data by user, these data cannot be subsetted together with the existing data in the atlases, and thus hinder any selective downstream analysis. As existing atlases are insufficient to allow subsetting of the growing data, scalable cloud web portals that can fulfil this property with performance and scalability are of urgent need ^21^. In the context of COVID-19 studies, such kinds of subsetting need to allow the quick retrieval of any combination - for example, age, ethnicity, gender, cell type, COVID-19 severity, and more - of input data for more refined downstream analysis.

Here, we present Scope+, an open source, transferable and scalable software architecture for cell atlas portal that facilitates effective meta-analysis downstream analysis for single-cell data. Scope+ architecture is realised in Covidscope web portal where we gathered approximately 5 million (4,899,035) single PBMC cells from 20 studies globally (Supplementary Table 2) and offered the data in the cloud as a valuable community resource. Scope+ addresses the high-throughput data challenges associated with dynamic subsetting and complex querying of 12.7B matrix count and ∼5M meta information with three key innovations. These are (i) database optimization strategies to allow extremely fast (10∼6000x faster) query of the 12.7B matrix based on the 5M meta data; (ii) simultaneous rendering of 4.9 millions of meta information in the website via implementing fastest MongoDB ObjectId based pagination in seconds; and (iii) novel architecture design to support large-scale bioinformatics data filtering, and visualization tasks in the web portal.

## Results

### Scope+: an open source innovative software architecture for large-scale data quick access and meta-analysis at cellular resolution

The Covidscope portal is designed based on a five-layered web application architecture, Scope+ (Fig. 2), with presentation layer, application logic layer and data storage layer being the three primary layers. MongoDB is applied for large scRNA-Seq data persistence in the database storage layer; while Flask is applied in the application logic layer to store web logics; and Plotly is applied for interactive graphics generation in the presentation layer (see Methods and Supplementary Materials). Four main functions of the portal are: fast cell sorting, interactive visualisation on cell level, visualisation on sample level, and data download (Fig. 1). Cell sorting is an interactive table which provides fast and intuitive cell-level search with metadata on the scRNA-seq data (Fig. 4a). Users can filter cells interested by various attributes such as sample, age, health status, and predicted cell types. Interactive visualisation on cell level includes the UMAP (see Methods) plot of the filtered cells colored by the metadata (Fig. 4b), and the selected gene expression across cell types (Fig. 4c). We also include sample level feature visualisation including cell type proportions (Fig. 4d), marker gene expressions, and pathway scores (Supplementary Figure 2) to provide comprehensive knowledge of the sample level information. Users can download all the queried cell results with the 10x genomics format along with the metadata and the feature level data on the website.

**Figure 1:**
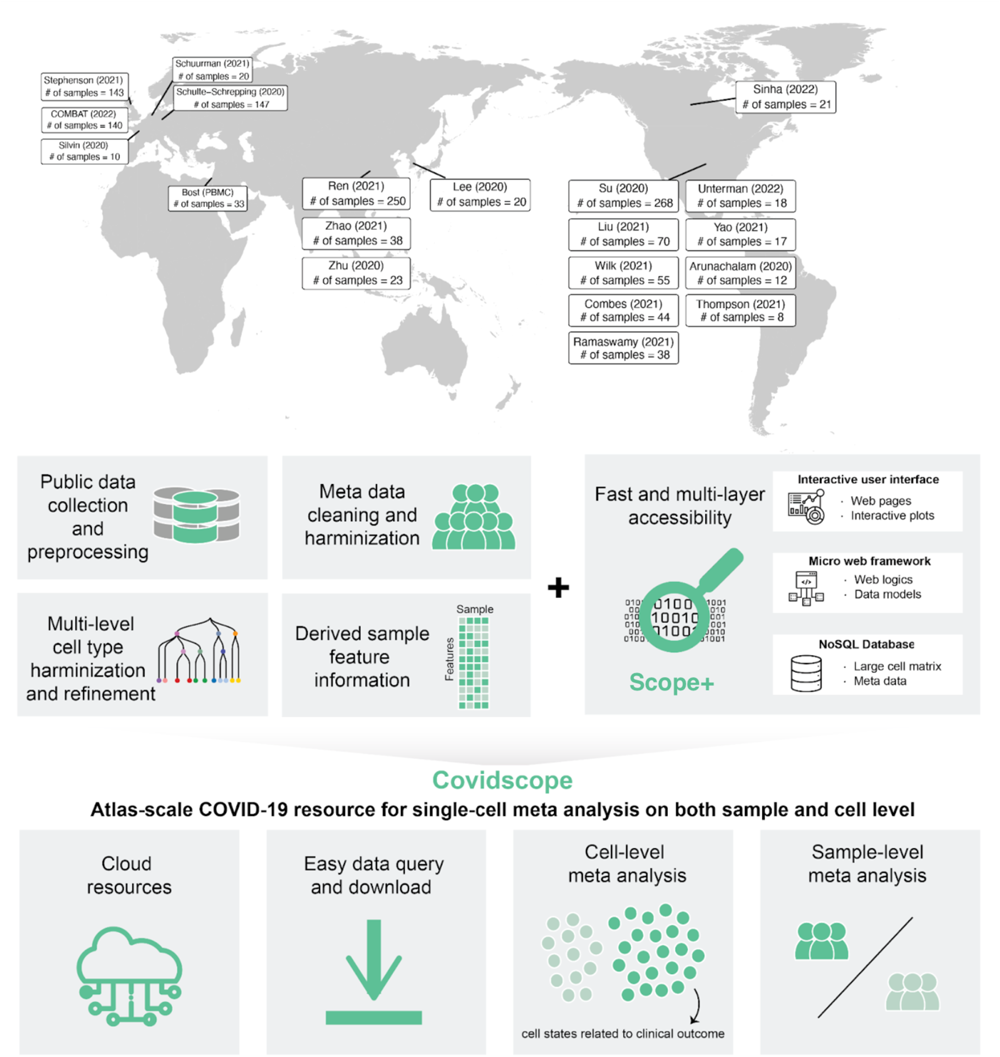
Schematic for large-scale COVID data integration, a portal in the cloud. Top panel shows the locations of the lead institutes of the 20 data sets. Bottom panel shows the schematic of the characteristics of the Covidscope web portal hosting the data sets and analytical results.

**Figure 2:**
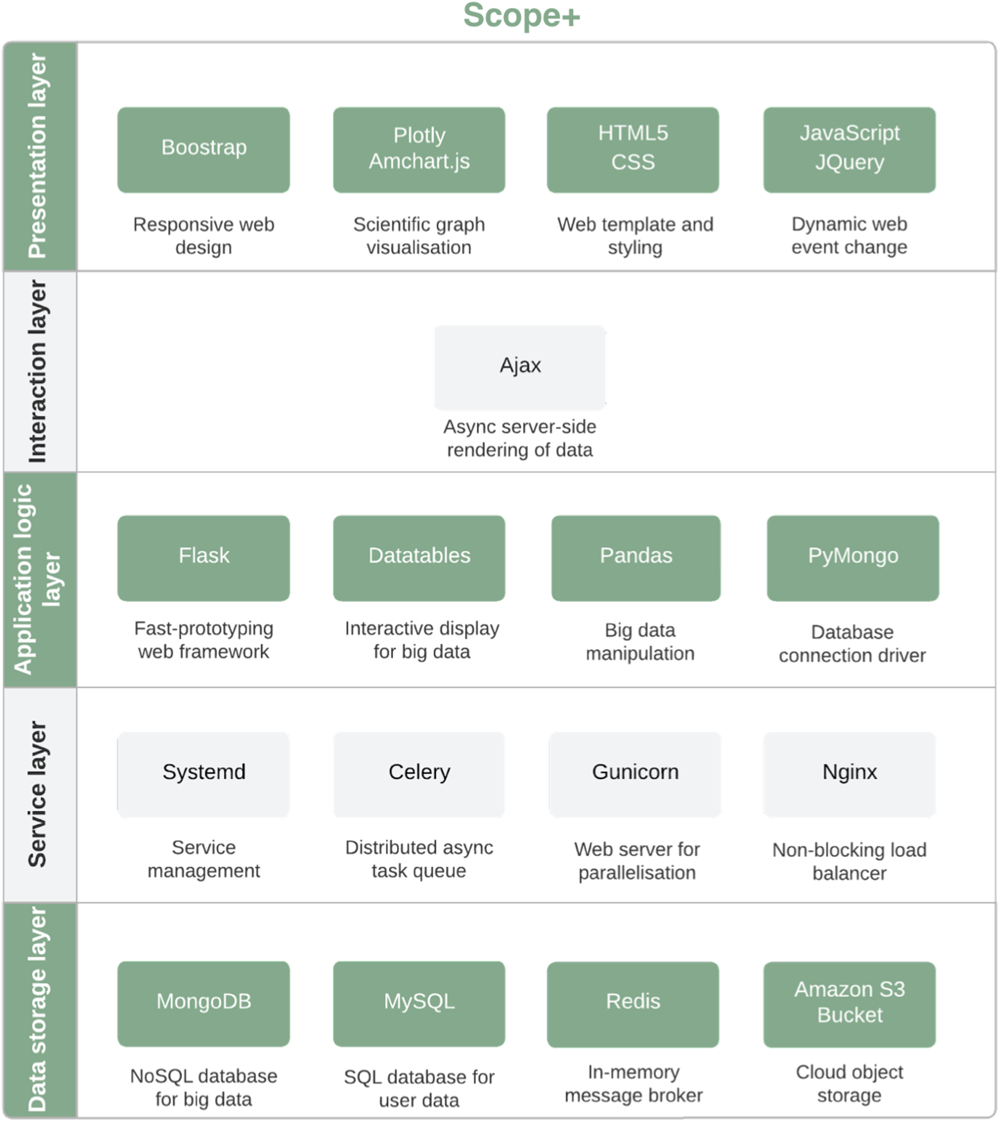
Scope+ architecture diagram behind Covidscope web portal. The Scope+ architecture behind Covidscope is divided into five layers i.e. presentation layer, interaction layer, service layer, application logic layer and data storage layer. Each layer is composed of one to several web application development tools with their functional description.

Thanks to our optimisation of the database using MongoDB indexing mechanism in Scope+, the performance of the cell sorting of the website is fast. We tested the performance of cell sorting by metadata and gene search in the UMAP plot across our optimised indexing and pagination strategies. The result is shown in Fig. 4e and f. The website queries cells by selecting condition through cell sorting within 0.06 millisecond; it can also search gene expression for selected cells from the large cell matrix (∼5 million cells and ∼30000 genes) within 6.7 seconds. It is by average >3 folds in cell sorting and >6000 folds faster than the baseline database. The result indicates that our noSQL based database can achieve fast query on single-cell omics data.

The atlas-scale data portal ready for single-cell meta analysis on both sample and cell level is achieved through two key components (Fig. 1). Firstly, a total of 20 clinical and study characteristics have been curated by reading through each paper for the relevant information and re-annotating each study to ensure consistency across the meta information (Table 2). This information enables the ease of subsequent meta-analysis across all data sets such as the analysis of a particular subgroup of patients.

Secondly, easy expression matrix accessibility through fast metadata subsetting is achieved by the modelling of the NoSQL database. The NoSQL database utilises the inherent dropout of the single-cell RNA-seq data, where only a small portion of the transcriptome is represented in each cell ^24,25^. It can index cells with cell identities featured by a series of sparse omics information and metadata as a key-values model. This solution reduces the data scale and enhances the data retrieval speed by indexing multiple keys. Particularly, we further enhance the performance of the website by three complementary optimisations including: (i) the choice of low level driver (ii) the pagination mechanism and (iii) the database indexing.

Our curated NoSQL database is primarily composed of four tables, these are (i) cell count matrix (ii) metadata of the cells, donors or data sets (iii) their computed UMAP coordinates and (iv) derived features for individual donors generated from the R package scFeatures ^26^. The design of our web application is highly modular, scalable, and optimised for runtime responsiveness. Together, this enables us to provide a centralised portal supporting quick database query, visualisation, data download, and integrative analysis of any type of single-cell transcriptomic data, not limiting to Covid-19.

### Covidscope: An COVID-19 single-cell portal

We applied Scope+ to our well-curated Covid-19 atlas collections of integrated data and released it in a web-based portal. Our integrated data collection consists of single-cell transcriptomics data from 20 published COVID-19 single-cell data sets (Table 1) that consist of close to 5 million cells in total from PBMC and/or whole blood extracted from almost 1,000 donors across 9 different countries using different protocols between 2020 and 2022. This portal is accessible at https://covidsc.d24h.hk.

**Table 1:**
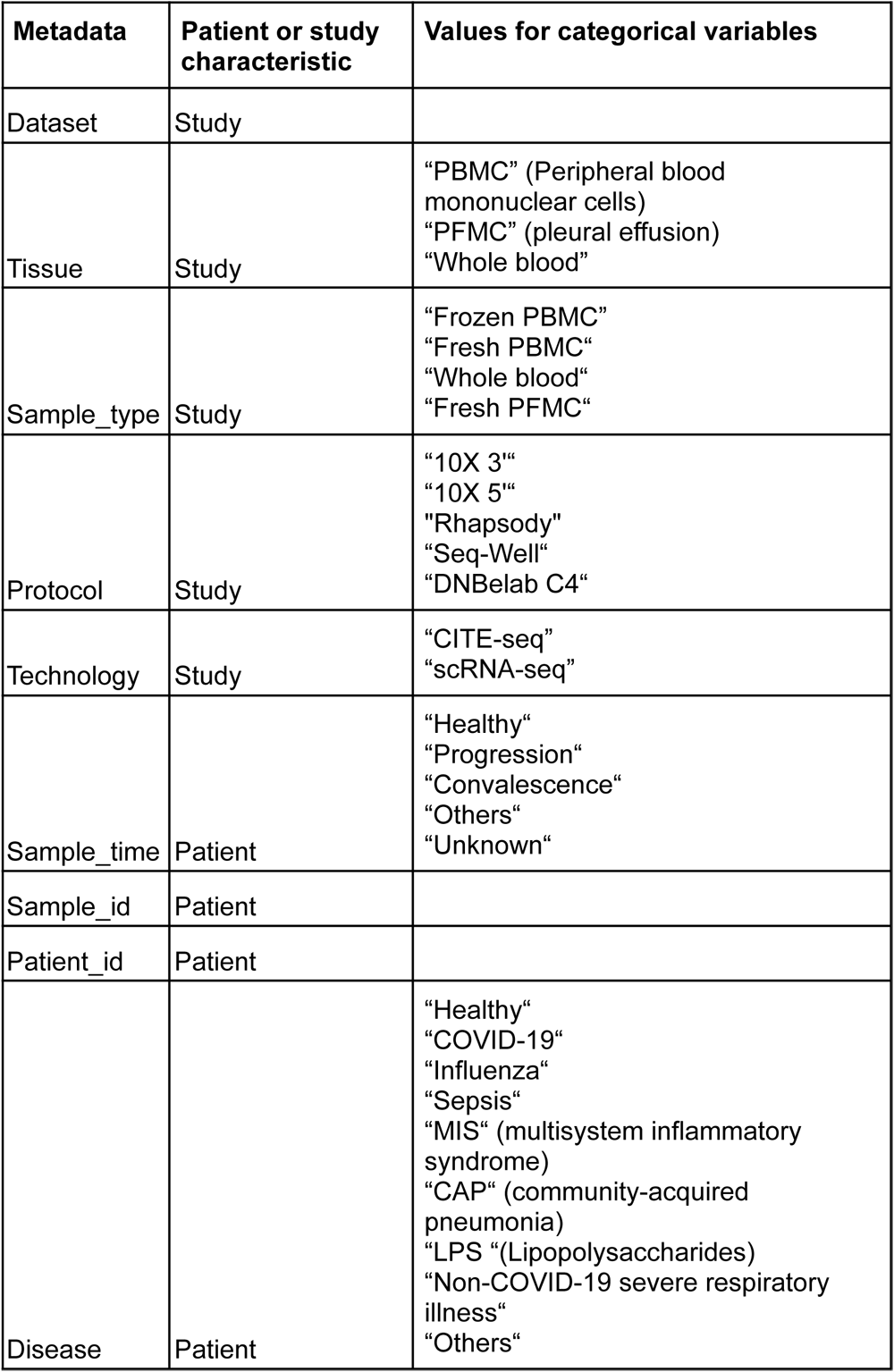
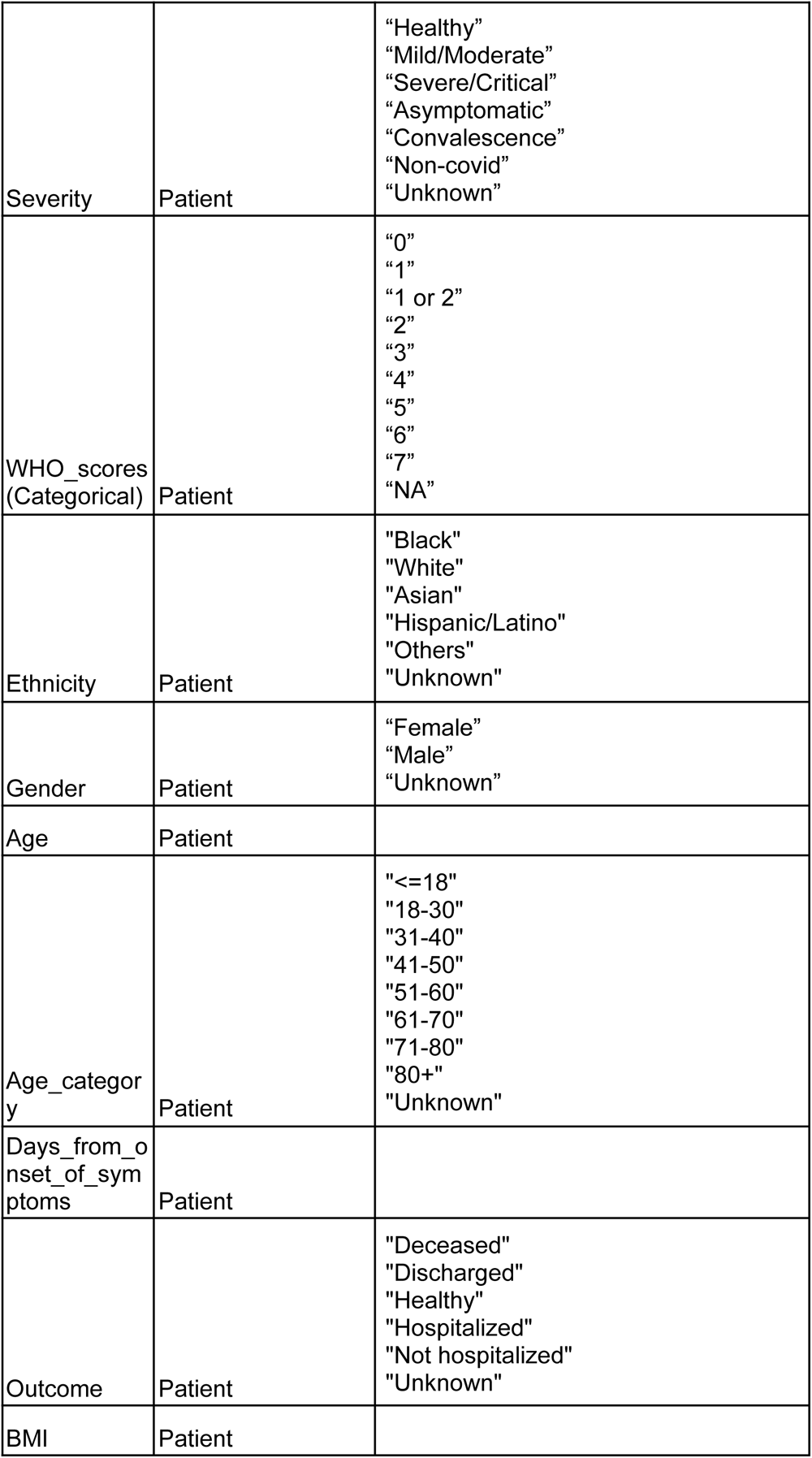
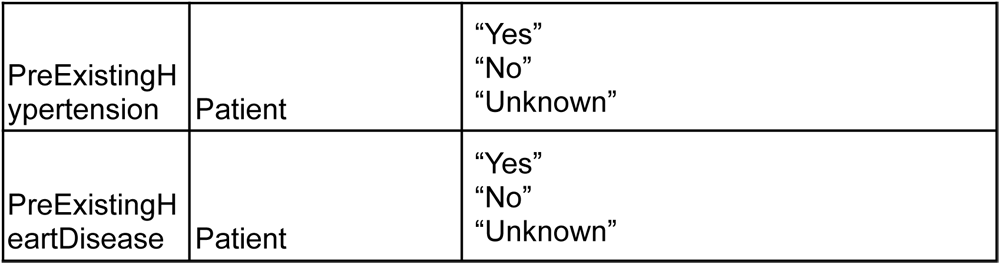
Overview of the metadata as curated in the 20 COVID-19 data sets. For categorical value types, we report the categories.

### Meta-features and derived features illustrate diversity in data collection

To construct a COVID-19 “analytic-ready cell atlas”, we downloaded the 20 published single-cell data sets (Supplementary Table 1) from public repositories and unified the technical, biological and clinical metadata. These data sets together contain close to 5 million cells in total from PBMC and/or whole blood using different protocols (10X, Seq-Well and DNBelab C4), published from July 2020 to March 2022. Together these data sets include 1,298 samples from 963 individuals, with diverse disease severity (15.2% Healthy, 40.8% Mild/Moderate, 32.8% Severe/Critical, 6.2% Non-COVID, 1.8% Asymptomatic, 0.5% Convalescence and 2.8% Unknown); disease outcome (39.4% Discharged, 7.4% Deceased, 6.4% Hospitalized, 1.0% Not hospitalized, 12.8% Healthy and 33.0% Unknown); gender (41.8% Female and 49.7% Male, 8.5% Unknown); age categories (2.6% <=18, 9.1% 18-30, 13.5% 31-40, 14.1% 41-50, 19.4% 51-60, 18.8% 61-70, 14.3% 71-80, 8.0% >80 and 0.2% Unknown) and sampling days from onset of symptoms (mean: 15.4 days, sd: 19.1 days) (Fig. 3a).

**Figure 3:**
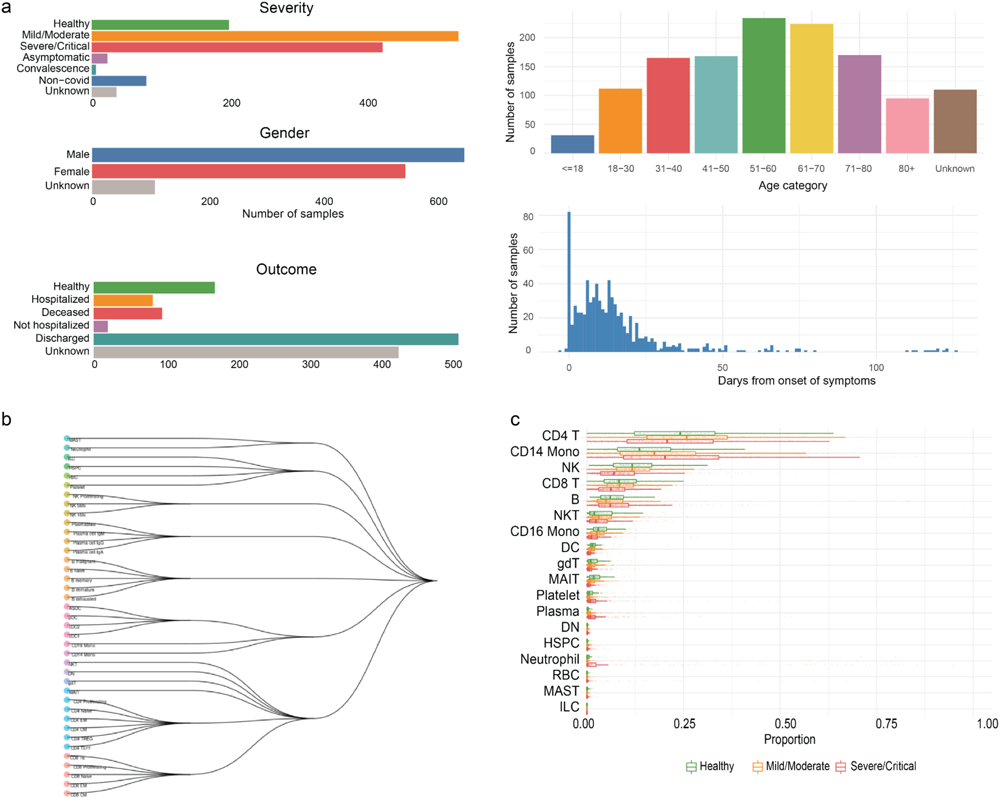
Data characteristics for the integrated COVID-19 data. a. Shows the distribution of selected patient characteristics. **b** Uses the HOPACH clustering algorithm to construct the hierarchy of the cell types in the COVID-19 data atlas. **c** Shows the proportion of the cell types in patients of different severity.

**Figure 4:**
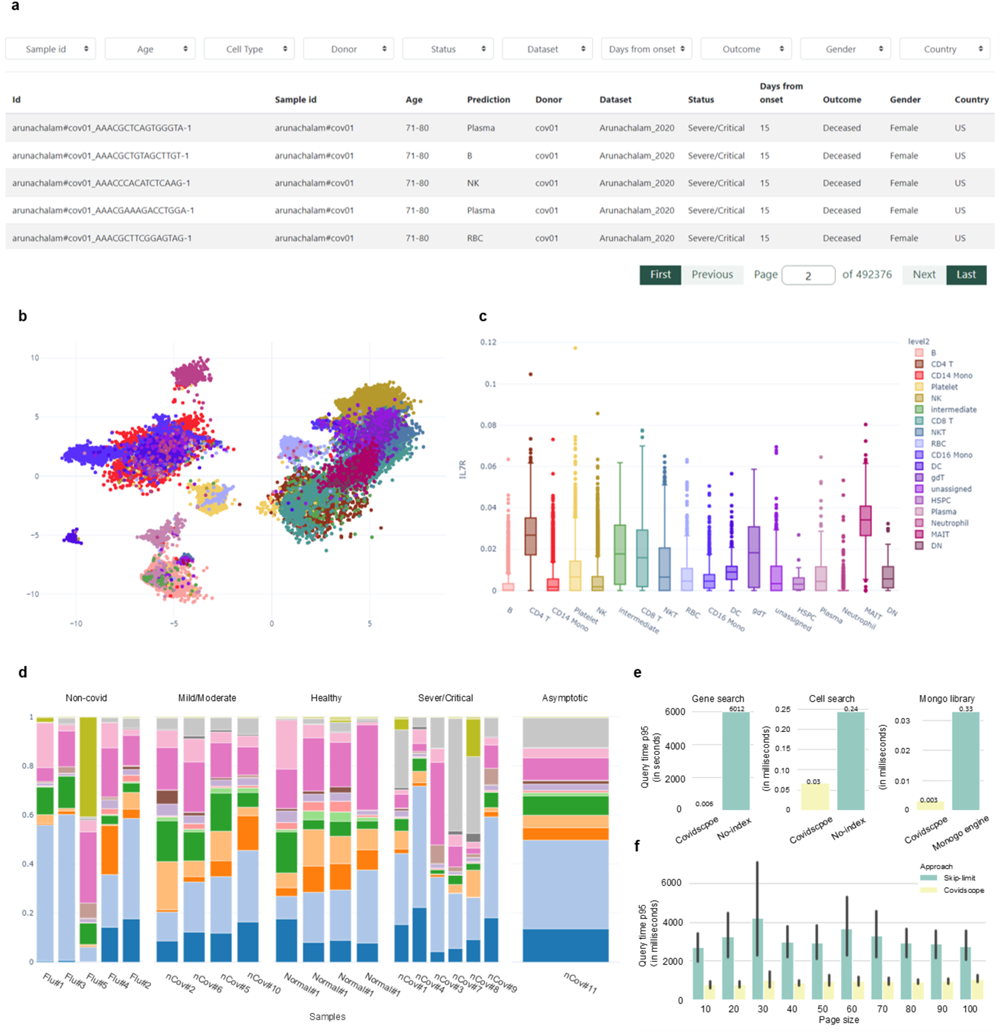
Covidscope web portal for COVID-19 single-cell meta analysis. **a** Demonstrates the interactive table for users to perform cell sorting through subsetting cells through metadata. Users can subset cells by attributes of interest. **b** Is the UMAP plot of cells from a specific data set (Lee et al, 2022) colored by the annotated cell types. **c** Shows the expression level of the selected gene across cell types. X-axis stands for the cell types while y-axis stands for the normalised expression level. **d** Is the cell type proportion across various disease severities of the Lee et al data set. Each bar represents one sample in the data set. Y-axis is the cell type proportion of each sample. **e** Demonstrates the time of different indexing test cases, each pair of the bar plots corresponds to gene expression searching, cell sorting, and library selection. **f** Demonstrates the time performance of pagination which x axis is the page size of the cell sorting table and y axis is the query time.

### Case study 1: cell level analysis of a subset cohort analysis

Integrated data from multiple studies with carefully curated metadata enables many new possibilities for downstream analyses. For example, using condition outcomes and cell type labels, one could perform case-control studies as well as multi-conditional studies examining composition change, expression shift, perturbation analysis and a range of other analyses in a cell type specific manner.

To demonstrate the power of an integrated data atlas, here we examined the existence of a common signature that distinguishes cells found in mild and severe COVID-19 patients across multiple data sets. We focused on the CD14 monocytes cell type as this is one of the key players in COVID-19. Using Leiden community detection algorithm, we obtained 23 subclusters in CD14 monocytes cells and identified a number of clusters that are significantly enriched in severe cells (Fig. 5a). In these clusters, the proportion of cells from severe COVID-19 patients reached over 75%, marginally higher than the overall proportion of “severe cells” in the entire CD14 monocytes population (53%), suggesting a potential “severe” signature. Of these clusters, cluster 18 potentially harboured common severe signatures across multiple cohorts, as it contained a significant number of cells (over 300) across seven data sets (Supplementary Figure 3a) and had a high data set diversity, as demonstrated by a high Shannon diversity index (Fig. 5b). Further identification of differentially expressed genes in cluster 18 compared to the remaining clusters (Fig. 5c) revealed that eight out of the top 10 up-regulated genes belonged to the immunoglobulin gene family (Supplementary Figure 3b). Multiple studies in literature consistently point to increased immunoglobulin antibodies level in severe patients compared to mild and healthy controls ^11,27,28^. Together, we demonstrate the data atlas promotes the identification of common severe signatures across multiple cohorts.

**Figure 5:**
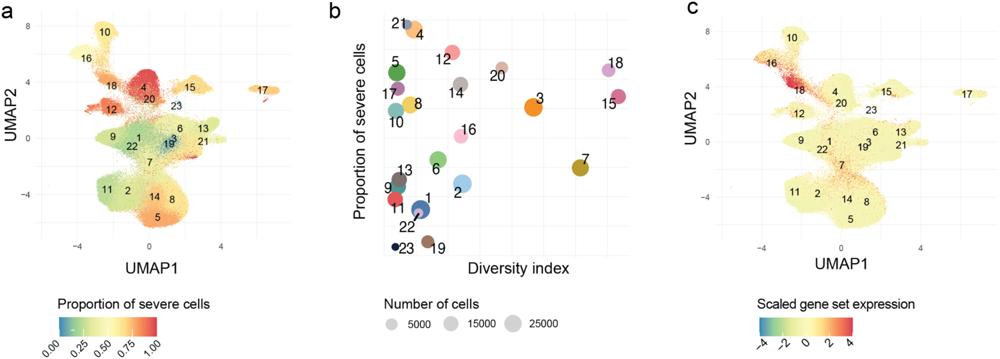
Case study on the CD14 monocyte cell population. **a** Shows the UMAP projection of the CD14 monocytes, with each derived cluster coloured by the proportion of severe cells in the cluster. **b** The diversity index quantifies the number of cells and number of data sets in each cluster, where a larger number represents more diversity. **c** The top 10 DE genes from cluster 18 and the remaining cluster were obtained. The scaled sum of gene expression from this gene set (IGLC3, IGLC2, IGHG2, IGHM, IGHG4, IGHG3, IGLC7, DEFA1B, FCER1A, IGHA2) were calculated for each cell and visualised.

### Case study 2: sample level analysis through integrative analysis of multiple data sets

One of the advantages of the cell atlas is that it enables the integrative analysis of multiple data sets, which opens the opportunity to examine various sub-populations across many data sets. One such question is the molecular difference underlying mild and severe outcomes in a given age group, which cannot be easily addressed without combining multiple data sets. For example, while ten studies in our atlas surveyed middle-aged patients of 41-50, the majority of these studies contain fewer than 10 mild patients and all studies contain fewer than eight severe patients (Supplementary Figure 4). Similar observations can be found in all other age groups, which pose a significant challenge for any downstream analysis. In contrast, combining all data sets enables us to examine 128 middle-aged patients of 41-50 and 131 elderly patients of 71-80 and derive biological discovery between mild and severe conditions in these two age groups (Fig. 6a).

**Figure 6:**
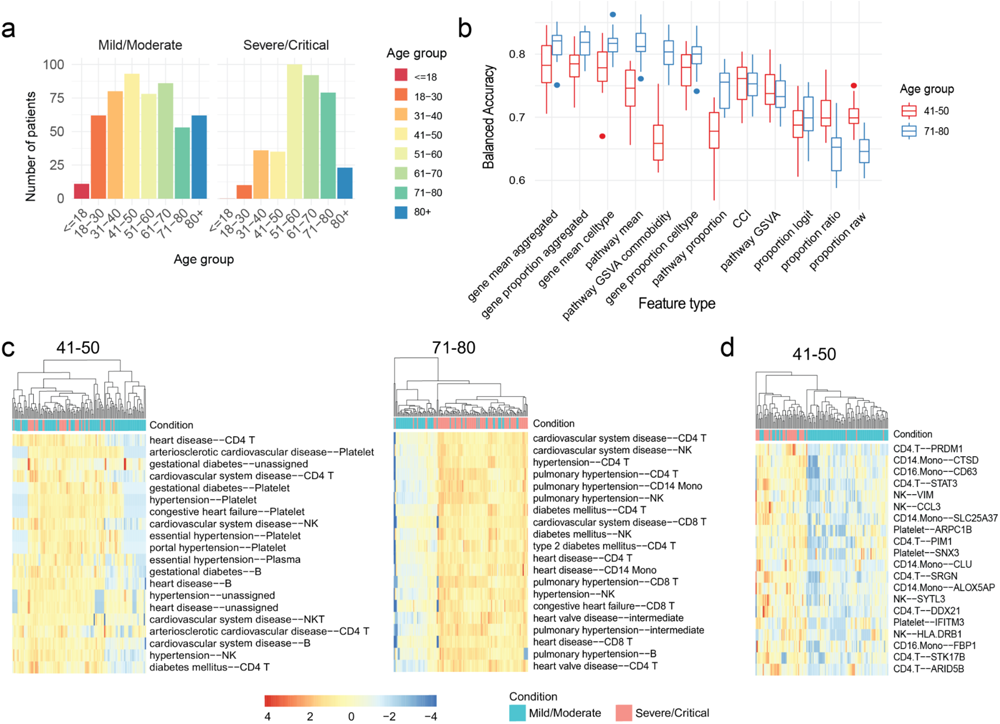
Case study of scFeatures on two distinct age groups of the COVID-19 patients. **a** Shows the age group distribution of mild/moderate and severe/critical patients in the database. In the study, we referred to all mild/moderate patients as mild, and all severe/critical as severe. We selected the 41-50 age group and the 71-80 age group and used scFeatures to generate molecular representation of individuals in each of the groups for further analysis. **b** Using mild and severe condition as the outcome, the boxplots show the classification accuracy using each feature type. Boxplots are coloured by age group. **c** Heatmaps show the values of the top 20 important features of the feature type “pathway GSVA comorbidity”, where the importance score was derived from classification model. Left shows the features on the 41-50 age group, right shows the features on the 71-80 age group. **d** Heatmap shows the values of the top 20 important features of the feature type “cell type gene mean” for the 41-50 age group.

To explore the differences between mild and severe patients, we used scFeatures to generate the molecular representation of each individual using a set of feature types. This was then followed by classification of the mild and severe conditions using each of the feature types. Majority of the feature types resulted in different prediction accuracy between the 71-80 age group compared to the 41-50 (Fig. 6b). For example, the feature type “pathway GSVA comorbidity”, which includes the pathway scores of several well-known comorbidities associated with COVID-19 outcomes ^29^ achieved an accuracy of over 0.8 for 71-80 but only over 0.65 for 41-50. Further examination revealed that the severe patients in the 71-80 age group had noticeably higher comorbidity pathway scores compared to the mild patients (Fig. 6c). This pattern was not observed in the 41-50 age group.

For the 41-50 age group, the top performing four feature types were all related to gene expression (i.e. “gene mean celltype”, “gene proportion celltype”, “gene mean aggregated” and “gene proportion aggregated”), achieving over 0.75 accuracy. Further analysis of top 20 features from the feature type “gene mean celltype” showed clear differential values between mild and severe patients (Fig. 6d). Additionally, we found that the cell type CD4 T, out of the total of 15 cell types, appeared 7 times out of the 20 features. This over-representation potentially suggests the importance of CD4 T cells in COVID-19 severity. Thus, through integrative analysis of patients across multiple data sets, we highlight the differing roles of certain feature types in affecting COVID-19 outcomes in different age groups and reveal biological insights into the mild and severe conditions.

## Discussion

In this paper, we propose an architecture Scope+ that enables fast access, query, and analysis of single-cell atlas at the scale of millions of cells. Using the architecture, we hosted a portal, Covidscope, containing almost 5 million PBMC from COVID-19 patients. Scope+ allows users to examine millions of PBMC cells, providing a doubling of data curation from the latest compilation at 2.2 millions ^21^ to 5 millions. The Covidscope portal provides users access at an unparalleled speed to COVID-19 gene expression data for cell clustering visualisation. Crucially, it enables flexible downstream meta-analysis by providing multi-conditional filtering and quick subsetting of a series of multi-view features including cell-cell interaction and pathway enrichment, thus providing deep insights into its pathogenesis and treatment.

Covidscope enables both exploratory and analytic data views. In the exploratory mode, users may engage with the original data by selecting certain cell types and studying the cells’ 2D projection. This is similar to many dashboard web portals. Here, we also present data under an analytical view where derived information is presented. For example, we generated a molecular representation of each individual using scFeatures, which enables users to examine the potential differences between moderate and severe patients by comparing their cell type proportions and gene expressions side by side. This feature allows researchers to quickly extract insights from the data without having to download hundreds of gigabytes of data and analyse it themselves. The portal is constructed to implement additional analysis features in the future depending on user requests.

Many COVID-19 scRNA-seq data will continue to be generated and published by groups around the world in the coming years. These data will differ in ways including (but not limited to) experimental conditions, patient characteristics, disease severity, and SARS-CoV-2 variants. Therefore, in addition to compiling and visualising all the existing data, our portal serves as a publicly available, easy-to-use tool for analysing both existing and new COVID-19 scRNA-seq data. In the future, users will be able to integrate new COVID-19 scRNA-seq with the existing data set of 5 million cells on our portal in a very fast, lightweight, and accurate manner. Tutorials for downstream analysis such as annotation, differential gene expression, trajectory inference, functional annotation, and cell-cell communication will be provided. The set of existing PBMC scRNA-seq data will be updated every few months. Users will have the option of whether or not to share their data. In addition, the current web portal is scaled to host 5 millions of PBMC data using our web stacks and configuration. The portal is ensured to accommodate further expansion of COVID-19 scRNA-seq data via the flexible design with: (i) NoSQL database model, (ii) fast pagination mechanism and (iii) database index. We can achieve scalability through either horizontal scaling (adding more machines) or vertical scaling (adding more compute resources).

Going forward, we will consider implementing more advanced features as requested by users into Scope+. For example, one advanced feature can be provision of an end-to-end pipeline from cell-annotation (scClassify), data harmonisation (scMerge2) and feature generation (scFeatures) for effective meta-analysis on user uploaded data. This will require not only providing resources on the cloud, but to further development of cloud computing components in our portal, allowing users to upload, store their own data and run the corresponding pipelines, respectively. In the current study, we have curated a comprehensive collection of COVID-19 data sequenced using scRNA-seq technology. With the explosion of multi-omics data sets, efforts are currently underway to expand our portal to include these multi-omics data.

The Scope+ architecture is one of the first generalizable atlas architectures made open source and is adoptable to users for reimplementation and a range of potential use cases. For extensibility, by open-sourcing our web portal, we invite researchers to extend or adapt our web portal to address their specific questions. We have made the web portal generalisable to support the hosting and exploration of any single-cell RNA-seq data. For citation and impact, we envisage that researchers who benefit from our framework will be likely to cite it in their work, therefore increasing the impact and citations of our work. For potential use cases, users can apply our architecture Scope+ to build such web portal based single-cell cell atlas that can visualize and analyze a selection of cells across all ranges of sources and be able to integrate and subset them by different criteria, e.g. cells from different atlases or datasets, different organisms, different species can be cross-compared at cell level, also users can select cells to hold-out for perturbational analyses and modelling.

In summary, as the availability of public large-scale single-cell studies continues to grow, our architecture Scope+ fills in the major gap in provision of a generalizable software architecture for cell atlas in terms of fast and flexible data access, harmonisation, query, and meta-analysis. A unique feature of Scope+ is that it enables provision of multi-level high-dimensional metadata subsetting and is transferable to any type of atlas-scale data for it to become readily available as a web portal. This distinguishing feature enables shared reproducible research in single-cell science. We envisage Scope+ will serve as an ever-growing “gold-standard” single-cell atlas portal architecture for the scientific community. Lastly, developing a cell atlas from scratch is time-consuming, Scope+ streamlines this process, which will significantly accelerate the atlas-scale single-cell research globally by making the cell atlases openly accessible into web portals for reproducible research.

## Methods

### Data Description

A collection of almost 5 million blood and immune cells extracted from almost 1,000 COVID-19 patients across 20 studies worldwide was used in this study. Supplementary Table 1 provides a summary of these 20 data sets, including the data accession ID. Each data set was transformed into a gene-by-cell raw count expression matrix. We performed size factor standardisation and log transformation matrix using the logNormCount function in the R package scater (version 1.16.2) and generated log-transformed gene expression matrices for downstream analysis. We performed cell type classification and re-annotated each data set with scClassify 23 for all data regardless of the original annotations. The data collection was merged and harmonised using scMerge2 (personal communication) in R software and stored in RDS file format.

### Data preprocessing

#### UMAP coordinates

UMAP coordinates were generated based on the top 50 principal components of the scMerge2 integrated matrix. The function *umap* implemented in the R package uwot was used, with *min_dist = 0.3*.

#### Patient based interpretable features

We applied scFeatures ^26^, a single-cell data feature extraction tool that we previously developed to extract a range of interpretable features. The features span across multiple categories, such as cell type proportion, cell type specific gene expression and cell type specific pathway enrichment and form the molecular representation of individuals. This provides molecular representations for all patients in the 20 COVID-19 studies, which are then visualised as interactive plots on the web portal.

#### Extract-transform-load (ETL) process

To process files for data import, Python scripts were used for converting RDS in section (i) to a CSV (comma-separated value) format, in which each line of the CSV file represents a record. The Unix command-line was used to import CSV to MongoDB using the “mongoimport” API. The imported data in the database follows the same structure as the original CSV, with a header as their column names. Data preprocessing scripts for the above mentioned processes is available on https://github.com/hiyin/scopeplus-scripts.

### Software architecture and implementation

The architecture Scope+ consists of three principal layers.

#### (i) Data storage layer

We designed separate collections for cell, cell type, and sample level features. Cell level features including the expression matrix, UMAP coordinates, and the cell type level feature as the metadata are stored in the **cov19atlas** database. The molecular features generated from scFeatures (see iv) including raw proportion, gene expression proportion and the pathway means score for each combination of sample and cell type are stored in the **scFeatures** database. More details can be found in Supplementary Figure 1. The molecular features generated from scFeatures (see iv) are stored into the **scFeatures** database. Single-cell meta information is stored into the **cov19atlas** database, with three collections namely single_cell_meta, umap and matrix, a universal id is created to cross-link these collections.

#### (ii) Presentation layer

HTML5 and Bootstrap (getboostrap.com) were used to build the modern responsive web user interface. JavaScript (www.javascript.com) and jQuery library (jquery.com) were used to manage dynamic content alteration, event, animations and asynchronous data transfer.

#### (iii) Application logic layer

A collection of APIs built on PyMongo, DataTables (https://datatables.net) and Flask make up the application’s logic. The *PyMongo API* handles the database connection and information retrieval. The self-constructed *search API* performs translation of search conditions into database query. The *DataTables API* displays paginated search results from MongoDB into tables using server-side rendering.

#### Data visualisation API for cell and individual level information

All diagnostic plots such as scatterplots, box plots, and bar charts presented on the website were implemented using the Plotly python package ^30^. On the cell level, we implemented UMAP plots to visualise the dimension-reduced cell expression using the *plotly.express.scatter* function. The box plot for gene expression levels across cell types was implemented using the *plotly.express.box* function.

We implemented several types of figures to demonstrate the individual level feature representation. The composition plot to visualise the cell type composition of patients was generated by the *plotly.express.bar* function. The visualisation of the distribution of individual feature values was implemented in several functions including box plot and heatmap. The box plot of pathway enrichment score across cell types was generated by the *plotly.express.box* function. The heatmap with dendrogram of mean gene expression and pathway enrichment score across cell types was generated by the *dash_bio* python package ^31^ using the function *dash_bio.Clustergram*.

### Optimisation

The Scope+ architecture implements three key optimisation strategies through:

1. Choice of driver: a low-level database driver PyMongo is used to interface the application with MongoDB. MongoDB cursor is returned directly from database query without converting back to Python object.
2. Pagination mechanism: MongoDB’s default ObjectId index is used for pagination of search results, server-side processing is enabled by setting “serverSide: true” in *DataTable API*.
3. Database indexing: frequently searched fields in MongoDB collections are indexed using *MongoDB API* (*db.matrix.createIndex({column_name:1}))*.

### Statistical analysis

#### Meta-analysis of CD14 monocyte cell populations

To demonstrate the power of using the atlas-scale database to reveal insights into COVID-19, we focused on the CD14 monocytes, which is one of the key cell types in COVID-19 response. Using *Monocle3* ^32^, which is found to be the best scRNA-seq clustering method for estimating the number of cell types ^33^, we clustered the CD14 monocytes into subpopulations. In order to prioritise clusters for downstream analysis, we used two criteria of (1) the proportion of severe cells in the cluster and (2) the diversity of the cluster in terms of the number of cells from each study and the number of studies that appeared in each cluster. We quantified criteria 2 using Shannon diversity index. Using these two criteria, we identified cluster 18 as a particularly interesting cluster as it was enriched in severe cells and had a high diversity index, containing a significant number of cells from seven studies. The standard DE analysis was then performed using scran to examine the up-regulated genes in cluster 18 compared to the remaining clusters.

#### Meta-analysis for individuals between 41-50 age group

To explore whether the mechanism underlying disease severity is the same across patients of different age groups, we selected the mild/moderate (referred to as mild) and severe/critical patients (referred to as severe) from two age groups of 41-50 and 71-80. We included all patients across all data sets in the atlas in order to obtain sufficient sample sizes. scFeatures was used to generate multiple feature types for each individual. To construct the comorbidity pathway features, we queried the Disease Ontology database ^34^ using the following terms of “heart”, “cardiovascular”, “hypertension”, and “diabetes” to obtain the relevant pathways and genes. Using the feature representations of individuals and the severity outcome, the prediction performance of each feature type was obtained using a linear kernel SVMwith three-fold cross-validation, repeated 20 times. As the outcome class has an imbalanced sample size, balanced accuracy was used as the evaluation metric.

### Hardware specification

The hardware specification of the hosting server is as follows; storage: 500BG boot disk and 2TB data disk storage; CPUs: 20 Intel(R) Xeon(R) Gold 6242R CPU @ 3.10GHz; memory: 64 GB RAM; operating system: Ubuntu 20.04.3 LTS (GNU/Linux 5.4.0-113-generic x86_64).

## Availability of source code and requirements

- Github link to software architecture Scope+: https://github.com/hiyin/scopeplus
- Portal home page: https://covidsc.d24h.hk (tutorial on portal https://covidsc.d24h.hk/tutorial)
- Operating system(s): Platform independent
- Programming language: Python
- License: MIT
- Any restrictions to use by non-academics: None

## Data availability

The web portal, data and meta-analysis are available on Covidscope (https://covidsc.d24h.hk/). User tutorials on how to implement Scope+ architecture with their atlases can be found at https://hiyin.github.io/scopeplus-user-tutorial/.

## Declarations

### Consent for publication

Not applicable.

## Funding

This work was supported by the AIR@innoHK programme of the Innovation and Technology Commission of Hong Kong to all authors, Research Training Program Tuition Fee Offset and Stipend Scholarship to YC. The funding source had no role in the study design; in the collection, analysis, and interpretation of data, in the writing of the manuscript, and in the decision to submit the manuscript for publication.

### Author contributions

JYHY and JWKH conceived and designed the study with input from YL and KHOY. DY, JC, and KHOY designed and implemented the database and web portal with input from YL, JYHY and JWKH. LM contributed to the user interface design and development for the web portal. YL and YC jointly curated the meta-data from the data collection. YC performed the case study analysis, implemented the data analytics and developed the corresponding R code with input from YL and JYHY. Everyone contributed to the generalisation and adoption of the portal’s software architecture on new data not in the portal. All authors wrote, read, reviewed the manuscript and approved the final version.

## Acknowledgements

The authors thank all their colleagues, particularly at D^2^4H, The University of Hong Kong, The University of Sydney, Sydney Precision Data Science Centre and Charles Perkins Centre for their support and intellectual engagement.

