## Supplementary Materials for "Scope+: An open source generalizable architecture for single-cell atlases at sample and cell levels"

#### Supplementary Tables

**Supplementary Table 1: Examples of published atlases that collate single-cell omics data from multiple studies and comparison to Scope+ / Covidscope.**

| Name | Publication | Web portal | Sample metadata / Analytical results / Both | Type of samples (human) | Number of single cells* (million) | Open source? | Generalizable architecture with source code | Integrative analysis at cell level |
| --- | --- | --- | --- | --- | --- | --- | --- | --- |
| Scope+ | / | https://covidsc.d24h.hk | Both | Healthy and diseased | 4.8 | Yes | Yes | Yes |
| Human Cell Atlas | Regev et al. 2017 | https://data.humancellatlas.org/ | Sample metadata | Healthy and diseased | 27.1 | No | No | No |
| hECA | Sijie et al. 2022 | http://eca.xglab.tech/ | Both | Healthy | 1.09 | No | No | Yes |
| Tabula Sapiens Consortium | Tabula Sapiens Consortium* et al. 2022 | https://tabula-sapiens-portal.ds.czbiohub.org/ | Analytical results | Healthy | 0.5 | No | No | No |
| HuBMAP | HuBMAP Consortium, 2019 | https://portal.hubmapconsortium.org/ | Sample metadata | Healthy | Not stated | Yes | Only web portal UI | No |
| Human Tumor Atlas Network | Rozenblatt-Rosen et al., 2020 | https://humanatlas.org/ | Both | Cancer | *Not stated* | No | No | No |
| Tumor Immune Cell Atlas | Nieto et al., 2021 | https://singlecellgenomics-cnag-crg.shinyapps.io/TICA/ | Both | Cancer | 0.5 | Yes, shiny app | No | No |
| Cancer Single-cell | Zeng et al., 2021 | https://ngdc.cncb.ac.cn/cancerscsm/index | Both | Cancer | 0.64 | No | No | No |

| Expressio<br>n Map |  |  |  |  |  |  |  |  |
| --- | --- | --- | --- | --- | --- | --- | --- | --- |
| Single-Cell Immunology Of SARS-CoV-2 Infection | Tian et al., 2022 | https://atlas.fredhutch.org/fredhutch/covid/ | Analytical results | COVID-19 | 3.2 | No | No | No |
| COVID-19 Cell Atlas | Sungnak et al., 2020 | www.covid19cellatlas.org | Analytical results | Healthy and COVID-19 | Not stated | No | No | No |
| SCovid | Qi et al., 2022 | http://bio-annotation.cn/scovid/#/ | Both | COVID-19 | 1 | No | No | No |
| ToppCell | Jin et al., 2021 | http://toppcell.cchmc.org/ | Analytical results | COVID-19, cancer, and healthy | 0.48 (COVID-19) | No | No | No |
| DISCO | Li et al., 2022 | https://www.immuninglecell.org/ | Both | 15 tissues, COVID-19 | 2.6 (COVID-19) | No | No | No |
| TIGER | Chen et al., 2022 | http://tiger.canceromics.org/#/ | Both | Cancer | 2.1 | No | No | No |
| ABC | Gao et al., 2023 | http://abc.sklehabc.com | Both | Blood disease | Not stated | No | No | No |
| PlaqView | Ma et al., 2022 | https://www.plaqview.com/ | Both | Cardiovascular disease | 1.7 | Yes, shiny app | No | No |
| SPICA | Andreatta et al., 2022 | https://spica.unil.ch | Both | Healthy and disease | Not stated | No | No | No |
| TCAC | Zhou et al., 2022 | https://tacalerner.ccf.org/ | Both | Alzheimer's disease | 1.1 | No | No | No |
| CellDepot | Lin et al., 2022 | http://celldpot.bxgenomics.com | Both | Healthy and diseased | Not stated | Yes | No | No |
| scAPAtlas | Yang et al. 2022 | http://www.bioailab.com:3838/scAPAtlas/ | Both | Healthy tissues | 0.8 | No | No | No |
| IAAA | Shen et al. 2022 | http://galaxy.ustc.edu.cn/IAAA | Both | Autoimmune disease | 0.7 | No | No | No |
| Cellxgene Annotate | CZ CELLxGENE Discover | https://cellxgene.cziscience.com/ | Both | Both | 55.5 | Yes | No | No |
| scIBD | Nie et al. | http://scibd.cn/ | Both | Inflammatory bowel disease | 1.14 | No | No | No |

\*Number of single cells collated by the resource at the time of writing, September 2023

**Supplementary Table 2: Overview of the 20 COVID-19 PBMC data sets used in this study.**

| Data set | Data accession ID | Reference | Number of cells | Number of patients |
| --- | --- | --- | --- | --- |
| Arunachalam et al. | GSE155673 | <sup>36</sup> | 56,639 | 12 |
| Bost et al. | GSE157344 | <sup>37</sup> | 50,284 | 33 |
| Combat et al. | EGAS00001005493 | <sup>38</sup> | 783,704 | 140 |
| Combes et al. | GSE163668 | <sup>39</sup> | 111,990 | 44 |
| Lee et al. | GSE147507 | <sup>40</sup> | 59,572 | 17 |
| Liu et al. | GSE161918 | <sup>8</sup> | 411,902 | 47 |
| Ramaswamy et al. | GSE166489 | <sup>41</sup> | 271,267 | 32 |
| Ren et al. | GSE158055 | <sup>3</sup> | 999,462 | 151 |
| Schulte-schrepping et al. | EGAS00001004571 | <sup>9</sup> | 328,780 | 74 |
| Schuurman et al. | GSE164948 | <sup>42</sup> | 32,384 | 20 |
| Silvin et al. | E-MTAB-9221 | <sup>43</sup> | 6,960 | 10 |
| Sinha et al. | GSE157789 | 44 | 80,994 | 14 |
| Stephenson et al. | E-MTAB-10026 | 4 | 643,071 | 130 |
| Su et al. | E-MTAB-9357 | 45 | 538,210 | 143 |
| Thompson et al. | GSE166992 | 46 | 63,895 | 8 |
| Unterman et al. | GSE155224 | 47 | 80,789 | 10 |
| Wilk et al. | GSE174072 | 48 | 174,753 | 39 |
| Yao et al. | GSE154567 | 49 | 69,983 | 17 |
| Zhao et al. | CNP0001250 | 50 | 88,374 | 19 |
| Zhu et al. | CNP0001102 | 51 | 46,022 | 3 |
| Total |  |  | 4,899,035 | 963 |

### Supplementary Figures

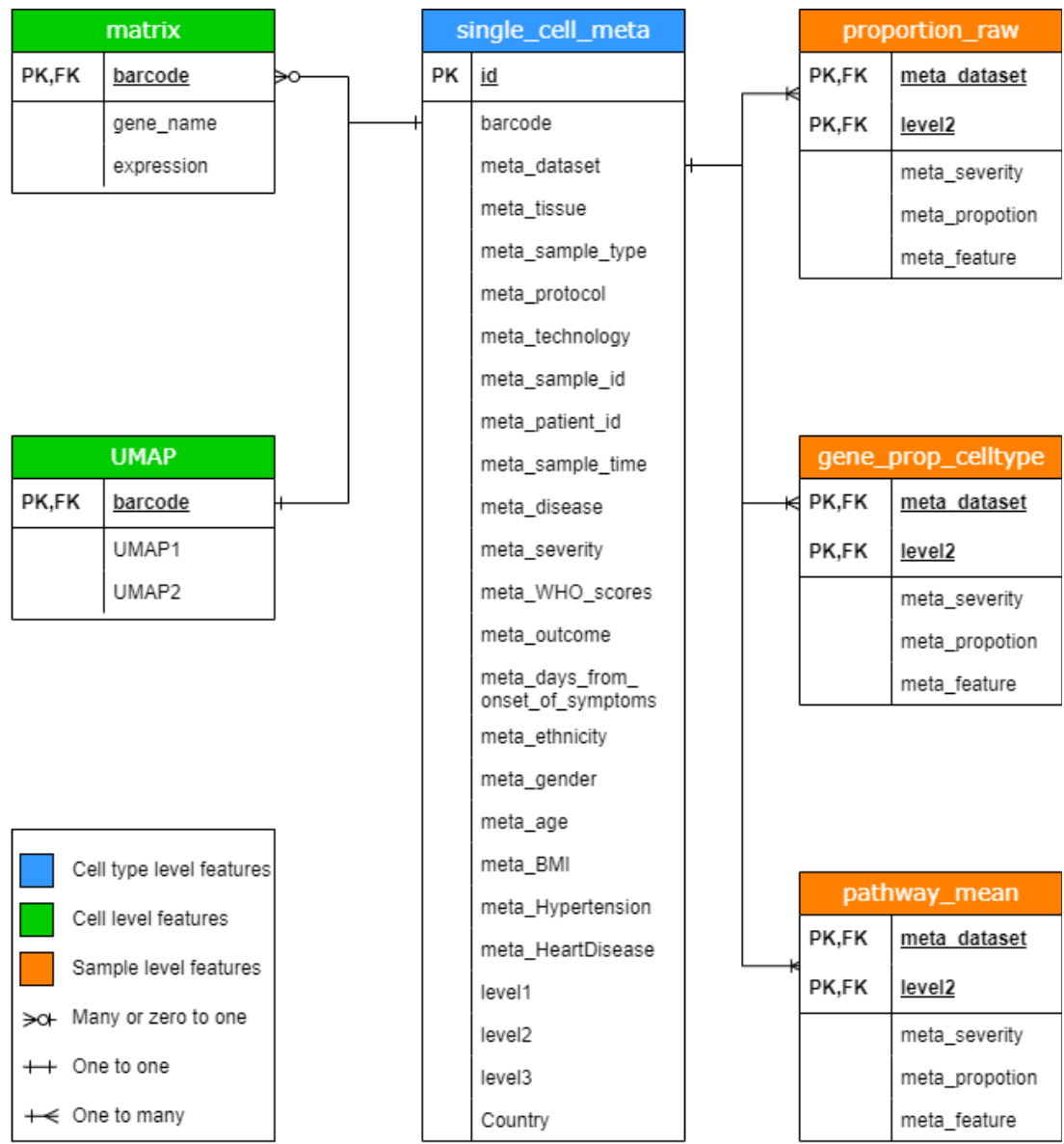

**Supplementary Figure 1. Database schema of the Covidscope data portal.**

Each rectangular box refers to an individual data collection in the MongoDB which Covidscope hosts. Primary keys (PK), foreign keys (FK), and other attributes within data collections are illustrated. Note that levels 1 ~3 refer to a different level of cell type annotation predicted by the scClassify. There are separate collections for cell, cell type, and sample level features. Cell level features include the expression matrix and the UMAP coordinates. The cell type level feature includes the metadata collection. Sample level features include the raw proportion, gene expression proportion and the pathway means score for each cell type. Cell and gene level collections are interconnected by cell barcodes. Sample level collections are connected by meta\_dataset and the level2 cell type annotation.

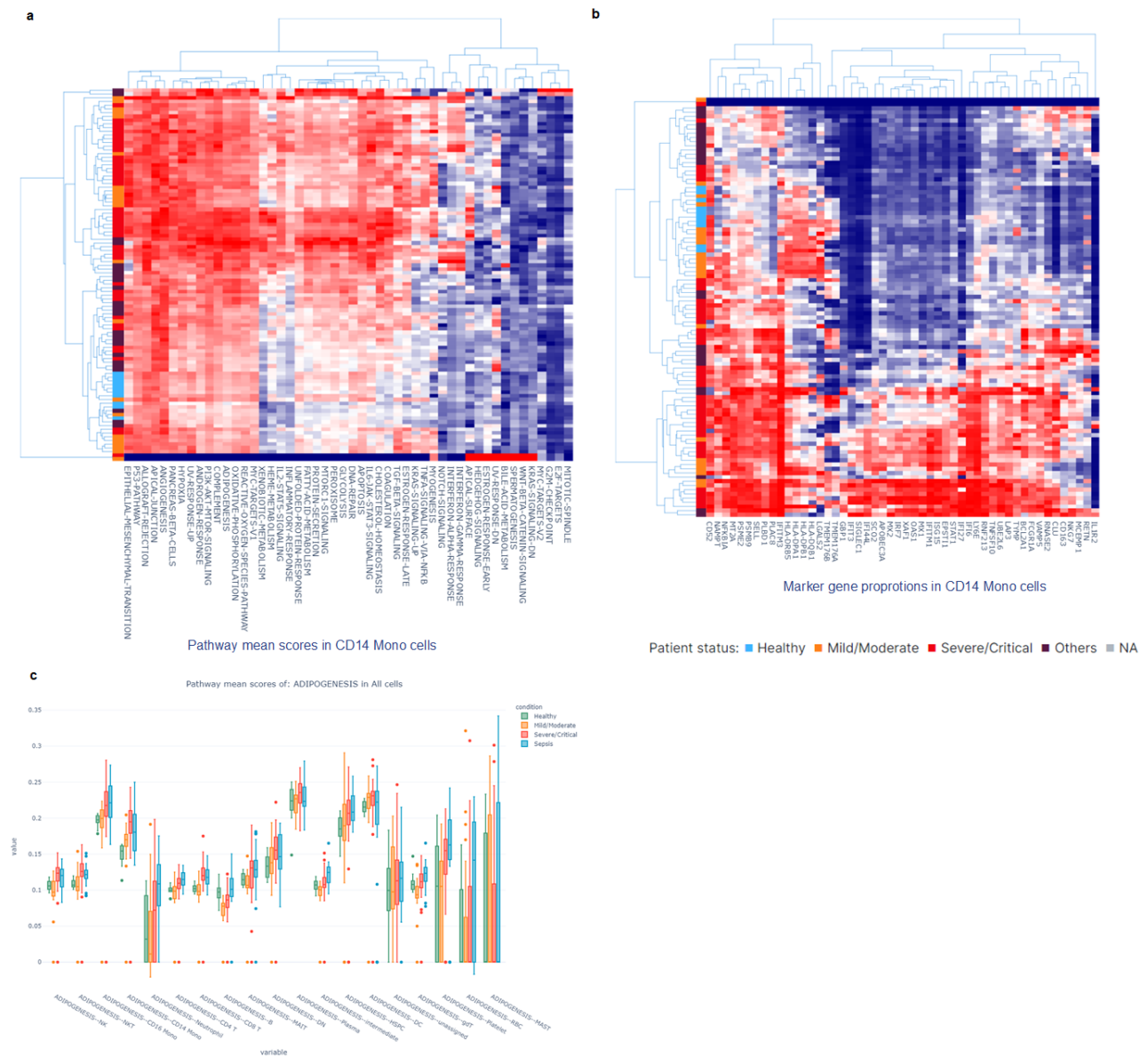

### Supplementary Figure 2. Screenshots for the sample level features on the websites.

**a** Heatmap shows the proportion of gene expression, constructed on cell type specific manner. **b** Heatmap shows the pathway score, constructed on cell type specific manner. **c** Boxplot shows the distribution of the pathway score, coloured by COVID-19 outcomes.

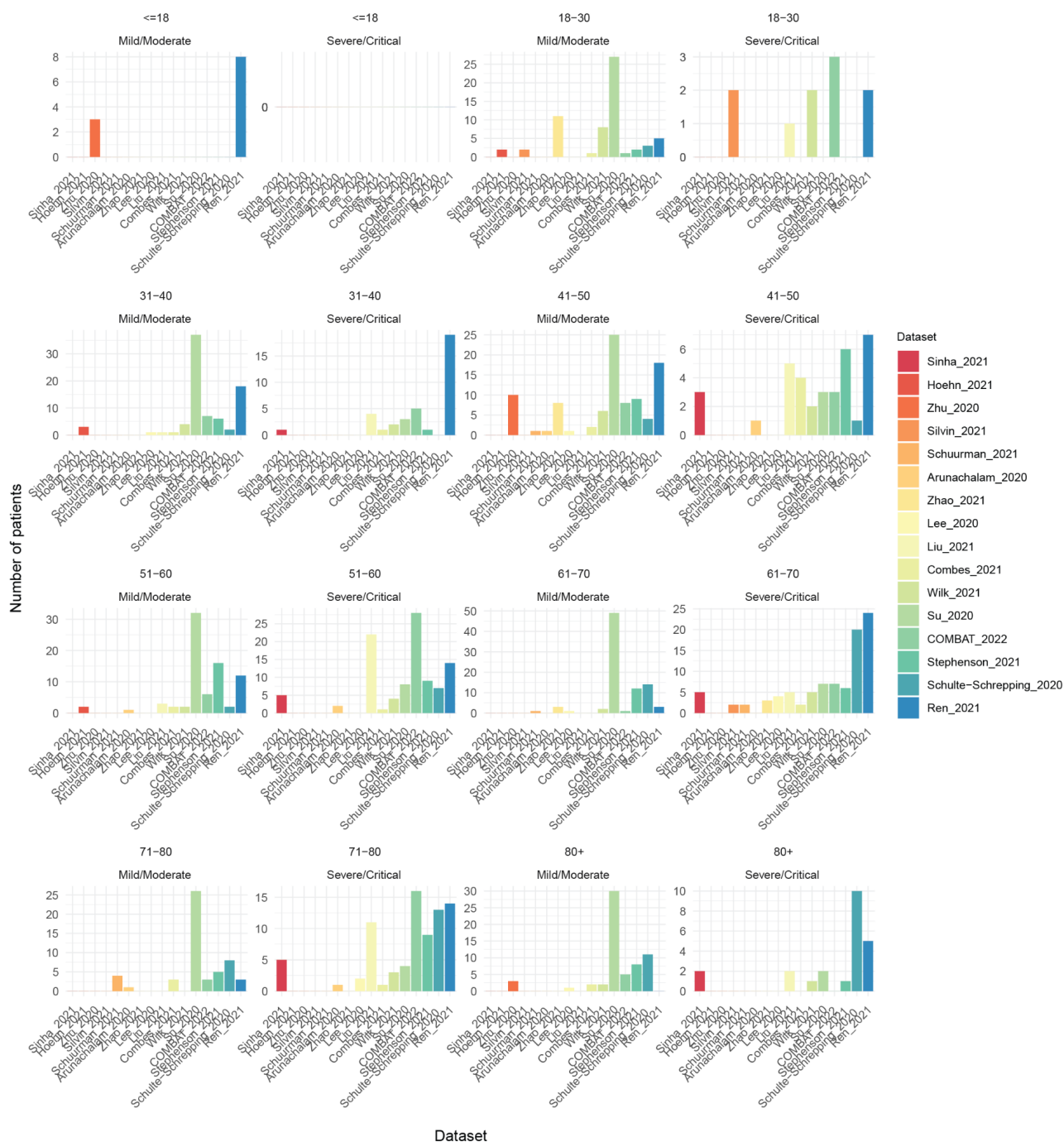

**Supplementary Figure 3. Age group distribution by data sets.**

Barplots show the age group distribution of the mild/moderate and severe/critical patients, coloured by data sets. Patients whose age is not reported in the original study were excluded. In the study, we referred to all mild/moderate patients as mild, and all severe/critical as severe.

**a**

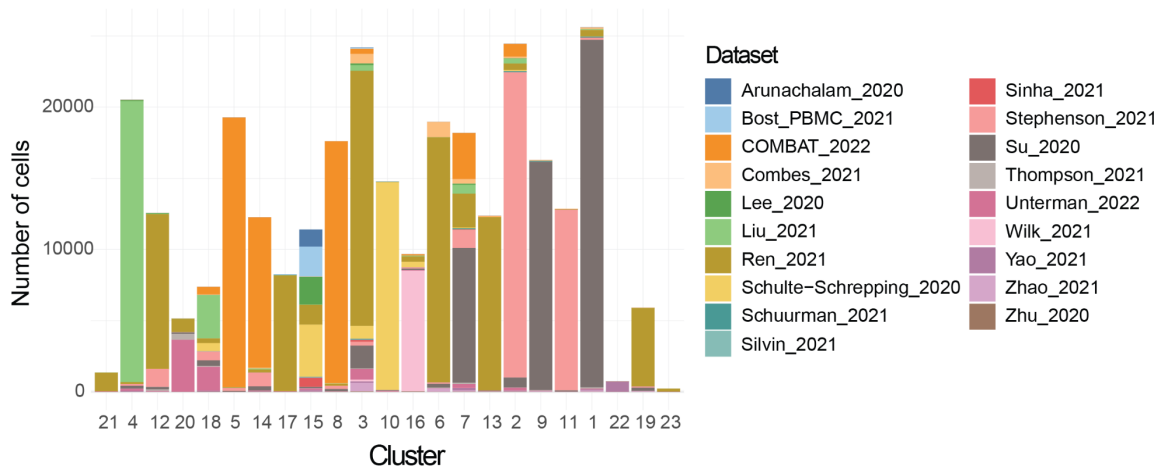

**b**

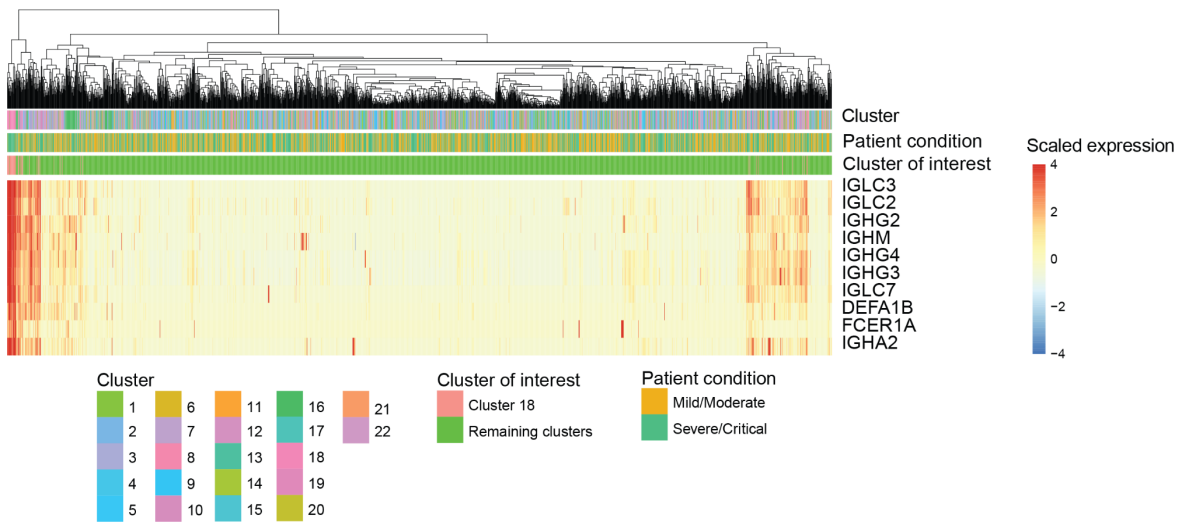

### Supplementary Figure 4. Analysis of the CD14 monocyte subclusters.

**a** The barchart plots the number of cells belonging to each study in each cluster, with the clusters ordered by the greatest percentage of severe cells to the least percentage. **b** Shows the expression heatmap of the top 10 DE genes between cluster 18 and the remaining clusters.
